# Reconstruction of septin higher-order nano-size structures in ovarian cancer cells uncover susceptibility to the septin-targeting small molecule UR214-9

**DOI:** 10.64898/2026.06.09.731048

**Authors:** Negar Khazan, Cameron WA Snyder, Nicole Dawney, Elizabeth LaMere, Srinivasan Ekambaram, Niloy A. Singh, Chandhana Ravi, Rajiv Snape, Hannah Aichelman, Elizabeth Pritchett, John Ashton, Thomas Kay, Myla Strawderman, Naohiro Yano, Dan T. Bergstralh, Gwenyth C. Eichfeld, Jeanne N. Hansen, Helge Ewers, Kyukwang Kim, Rachael B. Rowswell-Turner, Scott A Gerber, Erdem Tabdanov, Aurelie Bertin, Nikolay V Dokholyan, Richard G Moore, Rakesh K. Singh

## Abstract

In cancer cells, septins assemble into enigmatic higher-order structures of 300-700 nanometers, including long needle-like filaments, thick perinuclear rings, and cytoplasmic bundles or aggregates. The absence of genetic or pharmacological tools to recapitulate these architectures in-vitro has impeded mechanistic studies of their formation, function, and therapeutic targeting. Here, first, determining the overexpression of septin-2 in epithelial ovarian cancer (EOC) and its association with increased mortalities and dependencies, we select SKOV-3 ovarian cancer cells as a tractable model in which septin supramolecular assemblies can be recreated in-vitro and interrogated. This system shows that the forchlorfenuron (FCF) analog UR214-9 remodels septin architecture, converting co-expressed human septin octamers (SEPT2–SEPT6–SEPT7–SEPT9–SEPT9–SEPT7–SEPT6–SEPT2) into large cytoplasmic aggregates. In parallel, transiently expressed SEPT2 is reorganized into septin-rich noodle-like filaments, perinuclear rings, and web-like networks encircling the nucleus upon UR214-9 treatment. Mechanistically, UR214-9 disrupts the incorporation of SEPT2, SEPT7, and SEPT9 into canonical septin hetero-octamers, resulting in assembly-defective or imperfect oligomers that preferentially reorganize into these aberrant higher-order structures. This aggregation likely prevents septin-2 migration during interphase-to-cleavage furrow transition in NRK-49F-SEPT2-EGFP homozygous cells and impacts SKOV-3 cytokinesis, cell proliferation, adhesion and invasion and migration while sparing ceramide transport to the Golgi, preserving ER and cis-Golgi structure. These effects manifested in reduced growth of ovarian, endometrial and breast cancer xenografts without attracting significant off-target engagements per the global transcriptomic analysis of JIMT1 breast cancer and PANC-1 pancreatic cells. UR214-9 treated animals showed observable safety in animals. Thus, a tool to recreate aberrant septin structures and identification of septins as a druggable cytoskeletal target for ovarian, endometrial, breast and pancreatic cancer by perturbing their hetero-octamerization assembly is presented.

**Significance:** We provide a method to reconstruct the higher-order septin architecture observed in cancer cells, to study their assembly and functions. Intriguingly, cancer cells tolerate hetero-oligomeric septins lacking specific subunits, suggesting that compositionally deficient oligomers are not efficiently targeted for degradation, unlike unincorporated septin monomers in normal cells. This tolerance may enable accumulation of structurally aberrant septin complexes acquiring long-needles, rings or thick-aggregates in disease cells. We further show that septin oligomerization can be pharmacologically perturbed. By integrating structural, cellular, and energetic readouts using in-silico techniques, we establish a quantitative framework for septin-targeted modulation, generating UR214-9 as a new chemotype that disrupts septin oligomeric assembly via preventing incorporation of SEPT2/7/9, into canonical hetero-octamers, causes defects in cytokinesis, altered cell migration, viability, and remodels septin–actin architectures, ultimately impairing tumor cell growth. Thus, pharmacological targeting of septin assembly represents a tractable strategy to perturb septin-dependent cellular processes in cancer and neurodegenerative diseases with reported septin dysregulation.

## Introduction

Septins hetero-oligomerize to form filaments, bundles, and rings within cells, though their biological functions remain unclear^1–5^. Septin family of proteins participates in cytokinesis, cell migration, vesicular trafficking, exocytosis, cilia formation, and chromosomal dynamics^6–11^,. However, altered septin expressions are observed in malignancies including those of the pancreas, kidney, lung, colorectal, skin, brain, endometrium, ovarian, breast, and other organs^12–17^. Translocation of the mixed lineage leukemia (MLL) oncogene into a septin-2 and -6 gene locus occurs in acute myeloid leukemia (AML)^18^. Septin mutations have also been identified in tumors of the large intestine, skin, endometrium, and stomach^19^. Epigenetic alterations of septin-9 are the basis for the development of a blood test as an epigenetic marker for colon cancer^20^.

Furthermore, septins dysregulate growth factor receptors critical to cancer progression, including maintaining the abnormal persistence of ErbB2 at the plasma membrane of gastric cancer cells by reducing ubiquitylation and degradation^21^. Septins cluster and stabilize plasma membrane proteins^22–23^ with membrane-associated septin-9 inhibiting ubiquitin ligase functions, thereby reducing EGFR degradation^24^. Additionally, septins regulate actin remodeling during cell migration, metastasis, and invasion^25^. Depletion of septin-9 in metastatic cancer cells reverses epithelial-mesenchymal transition (EMT), reducing cell spreading, migration, and invasion, thereby demonstrating that targeting septin-9 could effectively control metastasis, the most lethal aspect of malignancies^26^.

Malignancies related functions summarized above make septins an important therapeutic target to control their multifarious roles in tumor growth, cytokinesis, exocytosis, migration, drug resistance and metastasis^27^. However, the dynamic nature of their higher-order oligomeric structures, intergroup dependencies and intergroup exchangeability, and the numerous subtypes including splice variants with shared or independent functions, present significant challenges in designing small molecule therapeutics against septins. It remains unclear whether a particular septin, a group of septin family members, or a specific septin-polymeric complex should be targeted to block septin orchestrated tumorigenesis.

Furthermore, chemotypes capable of interacting with septins, other than Forchlorfenuron (FCF)^28^ and REM127^29^, remain unavailable, creating a major bottleneck for medicinal chemistry efforts targeting septins. Compounding this challenge is the absence of robust screening assays for septin-targeting agents. It is not yet clear whether screening compound libraries using a septin GTPase assay will yield effective therapeutics, or whether assays that capture septin scaffolding or oligomerization functions, likely more relevant to their biological roles, will pave the way to optimal anti-septin candidates. Although FCF was the first compound shown to disrupt septin filament assembly^28^ and alter septin dynamics by inducing enlarged, stable polymers^29^, and despite demonstrating anticancer activity in endometrial cancer^30^, its substantial off-target liabilities^31^ and the impractically high doses needed to inhibit cancer-cell growth remain major impediments to preclinical or clinical development. Only REM127 has so far entered clinical trials for treatment of Alzheimer’s disease

In this study, we explore the dependence of ovarian cancer on septin-2 and examine how UR214-9^32–33^, mechanistically differentiating from FCF, and REM127 interferes with septin octamerization and spatial expressions and affects ovarian cancer proliferation, cytokinesis, lipid trafficking, migration, invasion and adhesion, each of which are linked to septins. We address off-target engagements of UR214-9 via global transcriptomic analysis of breast cancer cells exposed to treatment. We examined the pan-cancer treatment potential of UR214-9 by testing its efficacy against ovarian, endometrial, and breast malignancies. We also assessed whether targeting septins causes observable toxicities in healthy mice. Together, these studies provide the first systematic, medicinal chemistry-driven translation of septin architecture and biology, and the first preclinical evaluation of an anti-septin octamerization agent with therapeutic potential for ovarian, endometrial, breast, and pancreatic malignancies.

## Results

### Septin-2/7/9 are overexpressed in ovarian cancer tissues and predicts increased mortalities

Kaplan-Meier analysis of microarray data available at https://r2.amc.nl/ ^34–35^ showed that overexpression of septin-2/7/9 mRNA predicts increased mortalities among EOC patients (septin-2:p=0.041, septin-7:p=0.0041, and septin-9:p=0.029) (Figure-1A). Septin-2 and -9 are also overexpressed in malignant stroma that supports EOC (septin-2:*p*=7.24e^-04^; septin-9:*p*=9.57e^-011^) (Figure-1B). Immunohistochemical analysis of ovarian tumor array stained for septin-2 demonstrated increased expression in high grade stage-1 and stage-2 EOC tissues than normal ovary and benign cystadenoma (Figure-1C-left). Immunofluorescence analysis showed increased septin-2 expression in a high-grade EOC tissue than a normal ovarian and a benign ovarian tissue (Figure-1C-right). Previously we had shown increased septin-2 expression in EOC tissue sub-types than normal and benign ovarian counterparts^36^. Additionally, septin-2 enrichment predicts poor prognoses for patients diagnosed with breast cancer (*p*=0.063), lung cancer (*p*=0.002), liver cancer (*p*=0.00037), melanoma (*p*=0.018), renal cancer (*p*=0.0096) and the patients diagnosed with urothelial cancer (*p*=0.02), colorectal cancer (*p*=0.03) and malignancies of endometrium (*p*=0.04). Association of septin-2/7/-9 with increased mortalities in diverse malignancies are shown in Supplementary Figure 1.

**Figure 1:**
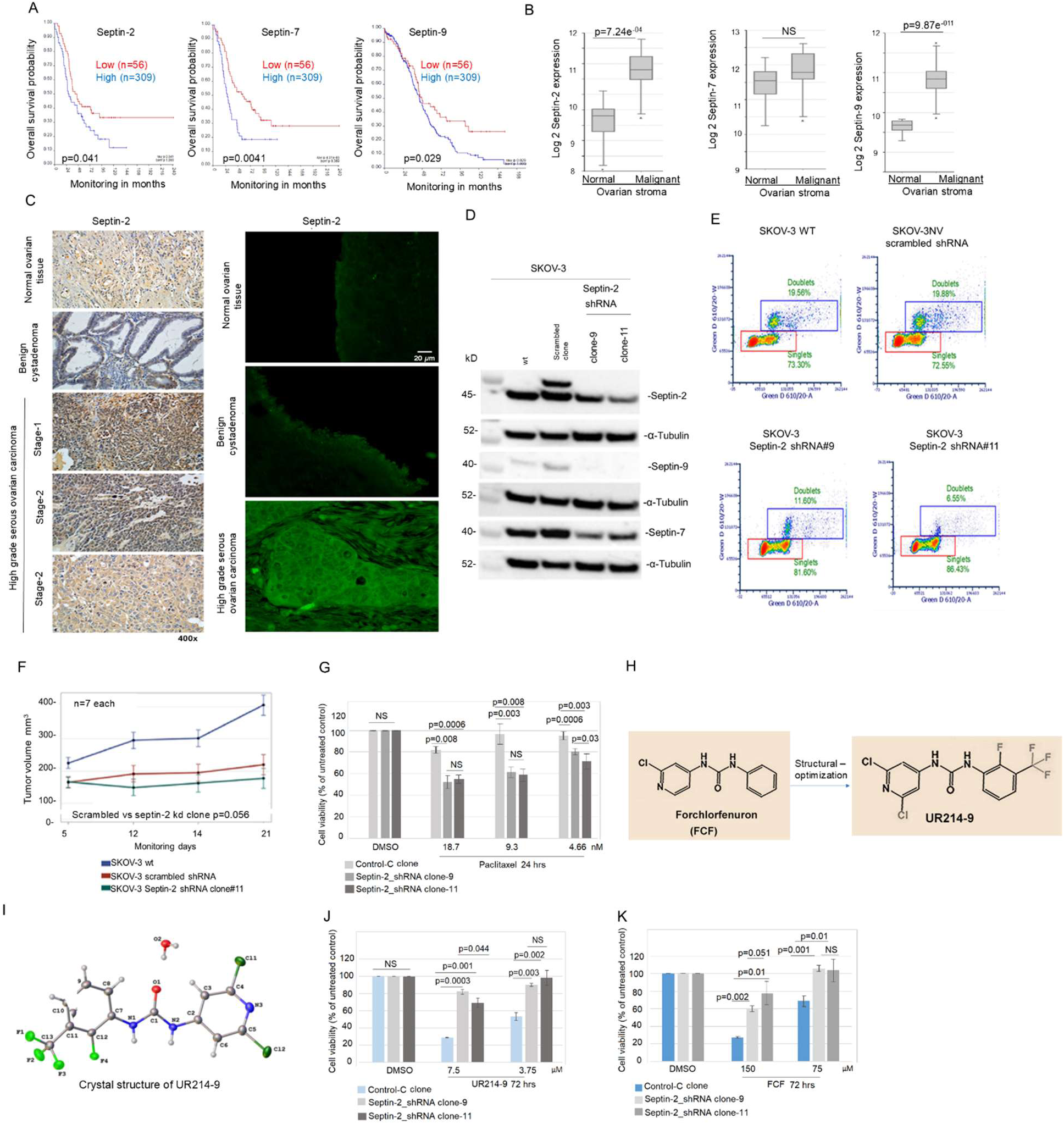
Septin-2 is a dependency factor for EOC. (**A**): Septin-2 mRNA overexpression predicts poor survival in EOC patients. For Septin-2, the Tumor Ovarian Adenocarcinoma database (Mechta-Grigoriou-107-MAS5.0-u133p2) database deposited at https://hgserver1.amc.nl/cgi-bin/r2/main.cgi was analyzed. (**B**): Expression of septin-2 in two stable knockdown clones #9 and #11 are shown. Stable knockdown of septin-2 completely knocked down expression of septin-9 in SKOV-3 cells. Septin –9 mRNA overexpression predicts poor survival in EOC patients. For septin-9 mRNA correlation with overall survival (OS), TCGA-381-tpm-gencode36 database deposited at https://hgserver1.amc.nl/cgi-bin/r2/main.cgi was analyzed. (**C**): Left: Immunohistochemical analysis of septin-2 expression in normal, benign cystadenoma, and high grade serous ovarian carcinoma (stage-1 and -2). Right: confocal analysis of septin-2 expression in normal ovarian tissue, benign cystadenoma and high-grade serous ovarian carcinoma tissues. Methods of staining and antibodies are described in Method section. (**D**): Partial stable septin-2 shRNA knockdown reduces septin-7 and -9 expression in SKOV-3 cells. (**E**): Partial septin-2/7/9 knockdown decreased doublet DNA and increased singlet DNA containing SKOV-3 cells population compared to wild-type and scrambled shRNA transfected cells. (**F**): Stable septin-2 shRNA knockdown clones formed slower growing tumors than the wild-type and null vector clones (p=0.056). (**G**): Stable septin-2 knockdown via shRNA increases response of SKOV-3 cells to paclitaxel in 24hrs. 1-tailed, paired T-test is shown. Similar increase in response to (−) epothilone-B was also observed (Supplementary Figure-3). (**H**): Chemical structures of FCF and UR214-9. UR214-9 is the optimized form of FCF. (**I**): x-ray structure of the purified UR214-9 is shown. The complete analytical co-ordinates of the x-ray crystallography are provided in Supplementary Figure-4. (**J**-**K**): Stable septin-2 knockdown via shRNA confers resistance to UR214-9 and FCF indicating dependencies of UR214-9 and FCF on septin-2/7/9 expression. Cell viability of UR214-9 and FCF treated cells increased in septin-2 partial knockdown clones than the scrambled shRNA clones during 48-72 hrs of treatment. 72-hrs data is shown. 1-tailed, paired T-test is shown.

### Septin-2 shRNA knockdown reduces Septin-7 and -9 expression, DNA-doublet population, tumor growth and enhances response to paclitaxel

The expressions of septin-2, -7, and -9 are interdependent^37–38^. We show that stable septin-2 knockdown, even partial, by septin-2 shRNA in SKOV-3 cells reduced expression of septin-9 and septin-7 (Figure-1D) along with septin-2 compared to scrambled control and wild-type cells. Uncropped gel images are shown in Supplementary Figure-2A. Next, we examined the impact of septin-2kd on cell division of SKOV-3 cells. Flow cytometric analyses showed that septin-2 kd clones (both 9 and -11) reduced DNA-doublet positive cells (11.8 and 6.56%) than >19% in SKOV-3 wild-type and scrambled control SKOV-3 cells (Figure-1E) indicating a modest G1-phase arrest^39–40^. Septin-2 knockdown resulted in a reduced rate of growth of tumors in NSG mice than SKOV-3 wt and scrambled shRNA when implanted in NSG subcutaneously (p=0.056, Figure-1F). Next, we investigated whether septin-2kd impacts the response to tubulin targeting therapies, paclitaxel and epothilones. Partial septin-2kD increases the responses of both paclitaxel (Figure-1G) and epothilone-B (Supplementary Figure-3) suggesting that septin-2 downregulation can enhance responses of EOC cells to paclitaxel and epothilone-B.

### Synthesis and x-ray crystallographic characterization of UR214-9. Structural optimization of FCF leading to UR214-9

(Figure-1H). Synthesis of UR214-9 was achieved as described earlier. The structure of UR214-9 was confirmed by NMR and Mass spectrometry. Purity was established by HPLC. Finally, structure was confirmed by X-ray crystallography. Detailed methods of x-ray crystallography of UR214-9 are described in Supplementary Information-4. UR214-9 forms a monoclinic colorless and block morphologic crystal (Figure-1I; Supplementary Figure-4). The asymmetric unit contains one molecule as a monohydrate in a general position. Hydrogen bonding (N-H…O, O-H…N, and O-H…O) links molecules in three dimensions. Molecules also appear to be aligned to maximize π-π interactions (∼3.3 Å).

### UR214-9 and FCF effects are septin-2/7/9 dependent

Next, we validated the interactions of UR214-9 and FCF with septin-2 using septin-2kd clones. We show that reducing septin-2/7/9 levels in SKOV-3 cells increases resistance to UR214-9 or FCF treatment (Figure-1J and K).

### UR214-9 binds greater than FCF but is inferior to natural ligands GTP and GDP

To understand the atomistic interactions of ligand binding in the septin family, we systematically compared the binding of GDP, GTP, and two small-molecule ligands (FCF and UR214-9) across septin 2, 6, 7, and 9 isoforms using molecular docking studies (Figure-2A). From the results, we elucidate that, despite sharing a conserved GTPase fold, these isoforms exhibit distinct ligand-binding properties (Figure-2B). We observed a pattern where GTP has a significant binding compared to GDP. For instance, septin9 showed the strongest GTP binding (–69.88 kcal/mol), followed by septin7 (–67.37 kcal/mol), septin6 (–60.73 kcal/mol), and septin2 (–62.74 kcal/mol). Whereas, other isoforms exhibited reduced GDP interactions, with energies ranging from –59.54 to –48.77 kcal/mol. Notably, septin9 maintained the strongest binding affinity for GDP (−59.54 kcal/mol) in the septin complex, while septin2 exhibited the weakest binding affinity with -48.77 kcal/mol. Notably, FCF and UR214-9 exhibited distinct binding affinities compared to GTP and GDP yet displayed clear isoform-specific binding. UR214-9 showed better binding than FCF across all septin isoforms. The strongest interaction for UR214-9 was observed in septin9 (−33.71 kcal/mol) and septin6 (−33.29 kcal/mol), suggesting these isoforms may possess allosteric pockets particularly receptive to the UR214-9 scaffold. FCF interactions were generally weaker, ranging from -26.65 kcal/mol (septin7) to -29.09 kcal/mol (septin6). These results suggest that UR214-9 may more effectively mimic key electrostatic or hydrogen-bonding features of nucleotide phosphate groups than FCF, enabling partial stabilization of the nucleotide-binding pocket. Overall, the results indicate that GTP has the greatest binding affinity, followed by GDP, UR214-9, and finally FCF, which could suggest a mechanistic basis for understanding the differential binding of GDP, GTP and small molecules (UR214-9 and FCF) to various septin isoforms.

**Figure 2:**
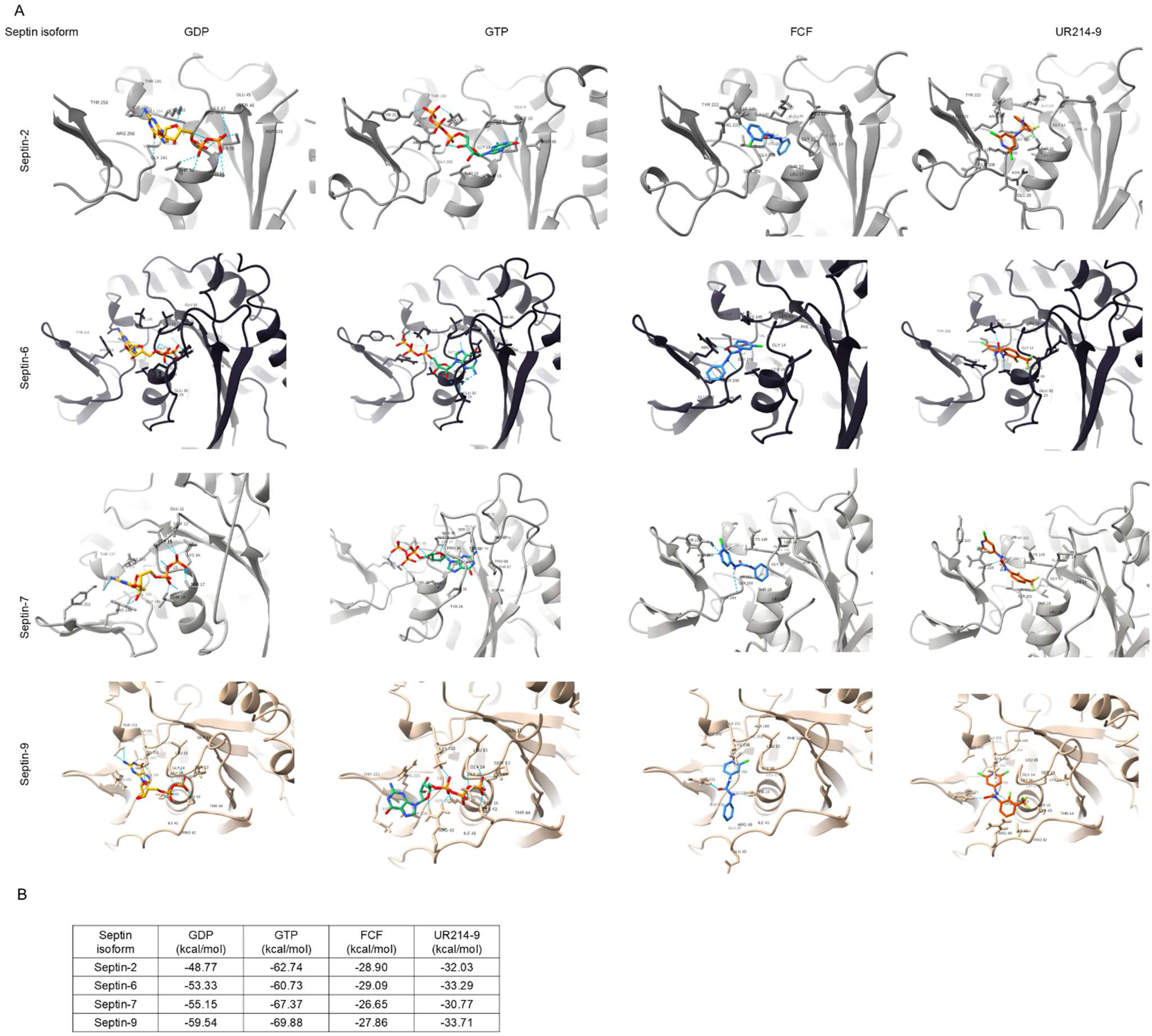
UR214-9 binds with septin isoforms differently. (**A**): Docking poses of GDP, GTP, FCF versus UR214-9 are shown. Interacting residues are shown. (**B**): Table shows the binding energies of GDP, GTP, FCF and UR214-9 against septin-2, 6, 7 and -9 in kcal/mol. It is intriguing that all the four ligands bind most tightly with septin-9 and least tightly with septin-2. This trend is similar to GDP which favors binding with septin-9 than septin-2.

### UR214-9 interacts better with the monomeric GTP-bound conformation of septin-2

Septin-2 binds GTP as a monomer and its dimerization is accompanied by GTP hydrolysis^40^. In the GDP-bound state, the switch region of both G-dimer subunits stabilizes the G-interface and forms closed conformer (Figure-2C). The closed conformer has strong intramolecular interactive forces (7.06A^0^) which stabilizes the dimeric complex that exhibits reduced affinity for a ligand. Switch residues Asp-101, Gly-104 that are in close interaction with GppNHp, are responsible for the homodimer formation stabilizing the septin-2 (inactive state). In GDP-bound state, in which septins are stable as dimer or oligomer, the GDP turnover is very low; and the phosphate groups of GDP masks all the P-loop residues that are required for binding with ligands such as FCF/UR214-9. Consequently, in the presence of GDP, UR214-9 are unable to increase the thermostability of SEPT2 dimer. On the other hand, the open conformer exhibits higher affinity for an external ligand due to the weak interactive forces (21.16 A^0^) with the switch residues (Figure-2D). The presence of γ-phosphate in GTP-bound state (active state) induces the conformational change of switch region and disrupts mutual interactions. GppNHp is a non-hydrolyzable GTP analog, and the GppNHp-bound Sept2 mimics the pre-dimeric GTP-bound state of sept-2. The GppNHp-bound structure carries ordered switch region, catalytic residues and has a Mg-ion coordinating with nucleotide. Having larger buried surface than the GDP-bound structure, GTP induces tighter binding between subunits in the GppNHp-bound structure. GTP bound-state, provides a less stable and open G-interface and institutes a higher GTP turnover rate. The Induced Fit-docking (IFD) shows that the interaction of ligand UR214-9 with the GTP-bound-like state of septin 2 (3FTQ) (score: -525.29) is better than with the GDP-bound state of septin 2 (2QNR) (score: -440.30) (Supplementary Figure-5).

### UR214-9 affects incorporation of septin-subtypes in septin polymers

Septins polymerize utilizing septin subunits from four different septin subunit families (2, 6, 7 and 9). We examined if UR214-9 or FCF treatment would affect oligomerization of septins. We treated SKOV-3. OVCAR-3 and OVCAR-8 cells with vehicle, UR214-9 3 µM and FCF 60 µM for 48 hours. We extracted proteins from the naïve/DMSO and treated cells in conditions that are shown to preserve oligomer^41^ and performed native gel electrophoresis^42^. As shown (Figure-3A-C), UR214-9 treatment reduced septin-2/7/9 levels in octamers in SKOV-3 cells. In OVCAR-3 and -8 cells, octamers were largely absent. We then examined whether septin-2 was potentially substituted by another septin-2 sub-group members (SEPT-1, 4, 5) in SKOV-3 cells. Re-probing the membrane showed that septin-4 did not substitute for loss of septin-2 in the SKOV-3 octamers (Supplementary Figure-2B). On the contrary, FCF did not interfere with septin-2/7/9 incorporation in SKOV-3 cells. Noticeably, FCF treatment seems to stabilize or concentrate septin-9 levels in octameric form in SKOV-3 cells. Neither agent interfered with hexamer formation. The Coomassie blue stained PVDF membrane is shown (Supplementary Figure-2B). Cartoon shows the sequence configurations of hexamers and octamers (Figure-3D). Next, examining the spatial arrangements of septins upon UR214-9, confocal microscopy showed that UR214-9 impacts the spatial distribution of septins in SKOV-3 cells. Treatment with UR214-9 (3µM) for 48 hours resulted in migration of septin-2 and -9 from perinuclear/cortex to cell membrane curvatures Figure-3E. Septin-6 was found to be reduced and septin-7 showed increased concentration around and inside the nucleus. Similarly, co-staining with septin-2/9/phalloidin-TRITC showed colocalization of septin-2/F-actin upon treatment with UR214-9 (Figure-3F). Notably, total septin-9 fluorescence/volume and total septin-9 fluorescence intensity showed increase compared to control (Figure-3Fright), whereas the nucleus volume and cell area were not altered indicating that cells were not facing cytotoxic effects. Septin-2 intensity did not change under treatment than vehicle (Figure-3Fright). Co-staining with phalloidin-TRITC to detect the impact on F-actin showed co-localization of septin-2 with F-Actin in OVCAR-8 and OVCAR-3 cells (Figure-3G-left). Treatment with UR214-9 caused nuclear accumulation of septin-7 and straightening of the curved septin-9 in SKOV-3 cells (Figure-3G-right).

**Figure 3:**
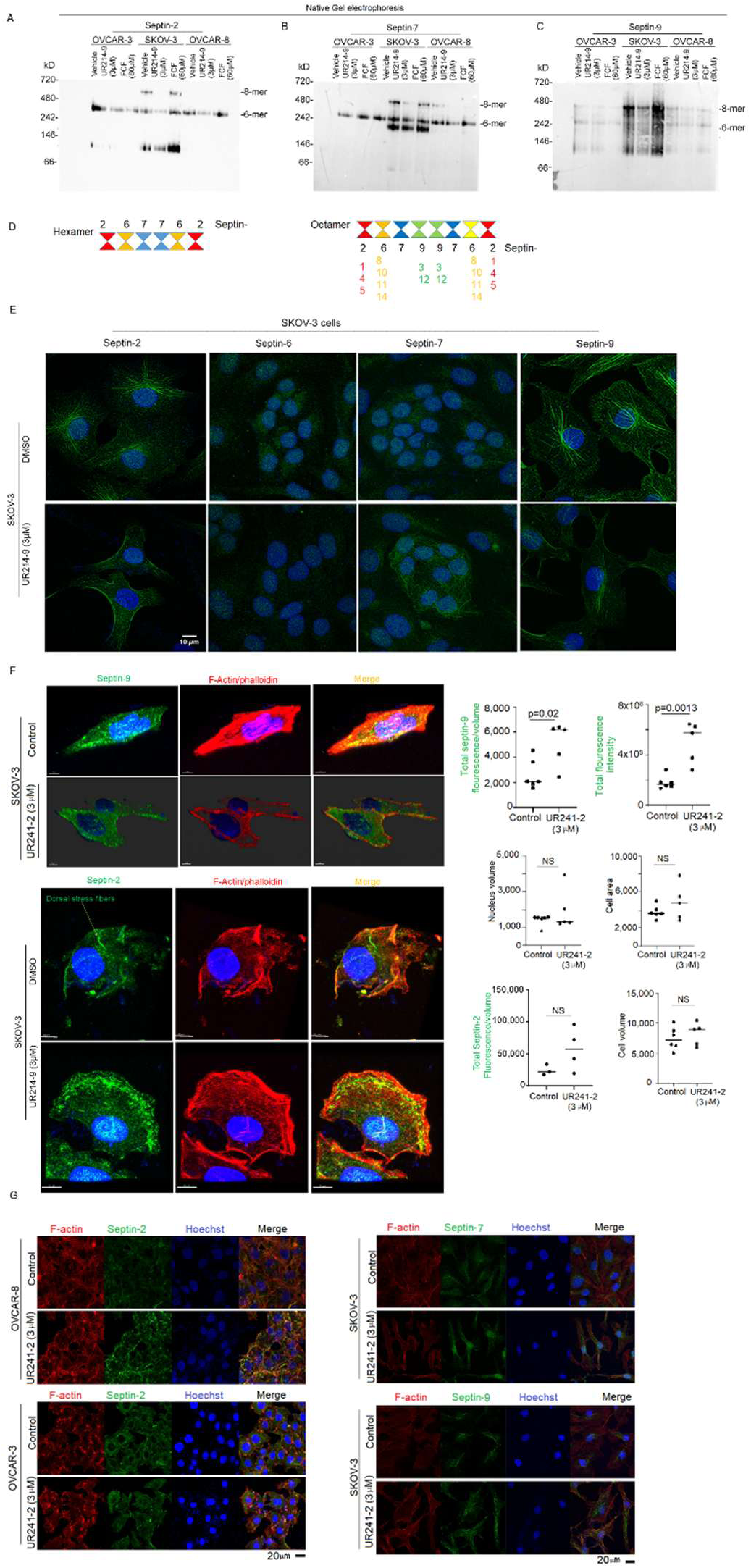
UR214-9 treatment inhibits incorporation of septin-2/7/9 subunit in hetero-octamer and alters spatial septin localization. (**A-C**): Native gel analysis of the OVCAR-3, SKOV-3 and OVCAR-8, cell-lysates treated with vehicle, UR214-9(3µM) or FCF (60µM) for 48 hours was examined to investigate the changes in polymer structures. UR214-9 suppresses septin-2/7/9 incorporations in octamers of SKOV-3 cells. Cells seeded in 100cc^3^ dishes were treated, washed, lysed using PIPES buffer+5mM NaCl solution and electrophoresed under native gel conditions using Coomassie blue at 210V for 1 hours, transferred overnight on PVDF membrane at 20V in cold room, treated transferred membranes with 10% acetic acid/MeOH, washed with MeOH to spot the ladder bands, images acquired using cell-phone cameras and stained with primary septin-2/6/7/9 antibodies and source matched secondary antibodies. Antibodies used are Septin-2/6/7/9 antibodies (Sigma Aldrich; septin-2: cat#: HPA018481; septin-7: cat#HPA029524; septin-9: cat#: HPA042564, 1:500). Notably, the octamers were not detected in OVCAR-3 and OVCAR-8 cells. (**D**): Cartoon depicts possible septin-hexamer and octamer constitution. (**E**): UR214-9 (3µM/48 hours) treatment induces migration of septin-2 and -9 from ventral stress fibers to curvature specific stress fibers in SKOV-3 cells but led to decreased septin-6 expression and strengthening of septin-7 filaments in SKOV-3 cells compared to control. Bar=10µm, 63×2 magnification. SKOV-3 cells (5,000 cells/well) seeded on EZslides overnight were treated with UR214-9 (3µM/48 hours), fixed with neutral paraformaldehyde (35µL) for 25 minutes (4°C), washed with PBST (3×500μL). Fixed cells were stained with septin-2 antibody (1:500, Sigma Aldrich; septin-2: cat#: HPA018481) overnight in PBST at overnight (4°C). Contents of the cells were removed, washed with PBST (3×500µL) and stained again with phalloidin solution (Vector Laboratories, 1:2000) overnight (4°C). Cells were washed (PBST, 5×500μL×10 minutes each). Well walls were dismantled, and glass slide was stained with Vectashield-DAPI solution and coverslip was applied and stored at 4 degrees under Aluminum file wrap. Cell images were recorded by the confocal microscopy. (**F**): Similar to (E), UR214-9 induced migration of septin-2 and -9 towards the cell’s curvature, interacting with F-Actin fibers. Septin-2 and septin-9 colocalize with F-Actin. SKOV-3 cells were seeded, treated, and stained with the Septin antibodies as described in section (E). F-actin was stained with Phalloidin-TRITC (ECM Bioscience, cat#PF7551; 1:1000 in PBST, overnight). Cells were washed (PBST, 5 times/30 mins and stained with DAPI containing mounting medium and confocal images were recorded. Total septin-2/cell volume and septin-9 fluorescence/cell-volume, total septin-9 fluorescence intensity, cell area, cell volume, nucleus volume in the SKOV-3 cells treated with DMSO/vehicle or UR214-9 was evaluated and calculated. Septin-9 fluorescence significantly increased in the treatment group. (**G**): (left) F-actin and Septin-2 co-localize in OVCAR-8 and OVCAR-3 cells. (right-upper): Septin-7 shows nuclear concentration in SKOV-3 cells upon treatment with UR214-9. (right-lower) Septin-9/F-actin also exhibit co-localization in SKOV-3 cells. SKOV-3, OVCAR-3 and OVCAR-8 cells were seeded in 8 well slide-chambers and treated, processed and stained with septin2/7/9 antibodies/corresponding secondary antibodies and Phalloidin-TRITC as described in (E). Confocal images were recorded.

### UR214-9 treatment forms large size aggregates of septin octamers

We next examined impact of UR214-9 on purified human octamers in vitro. Purified human octameric septins (Sept2-Sept6-Sept7-Sept9-Sept9-Sept7-Sept6-Sept2) upon incubated with UR214-9 at 10 µM for 30 minutes in a 50 mM Tris-HCl, pH 8, KCl 50 mM buffer showed large aggregates, much larger in sizes (Figure-4).

**Figure 4:**
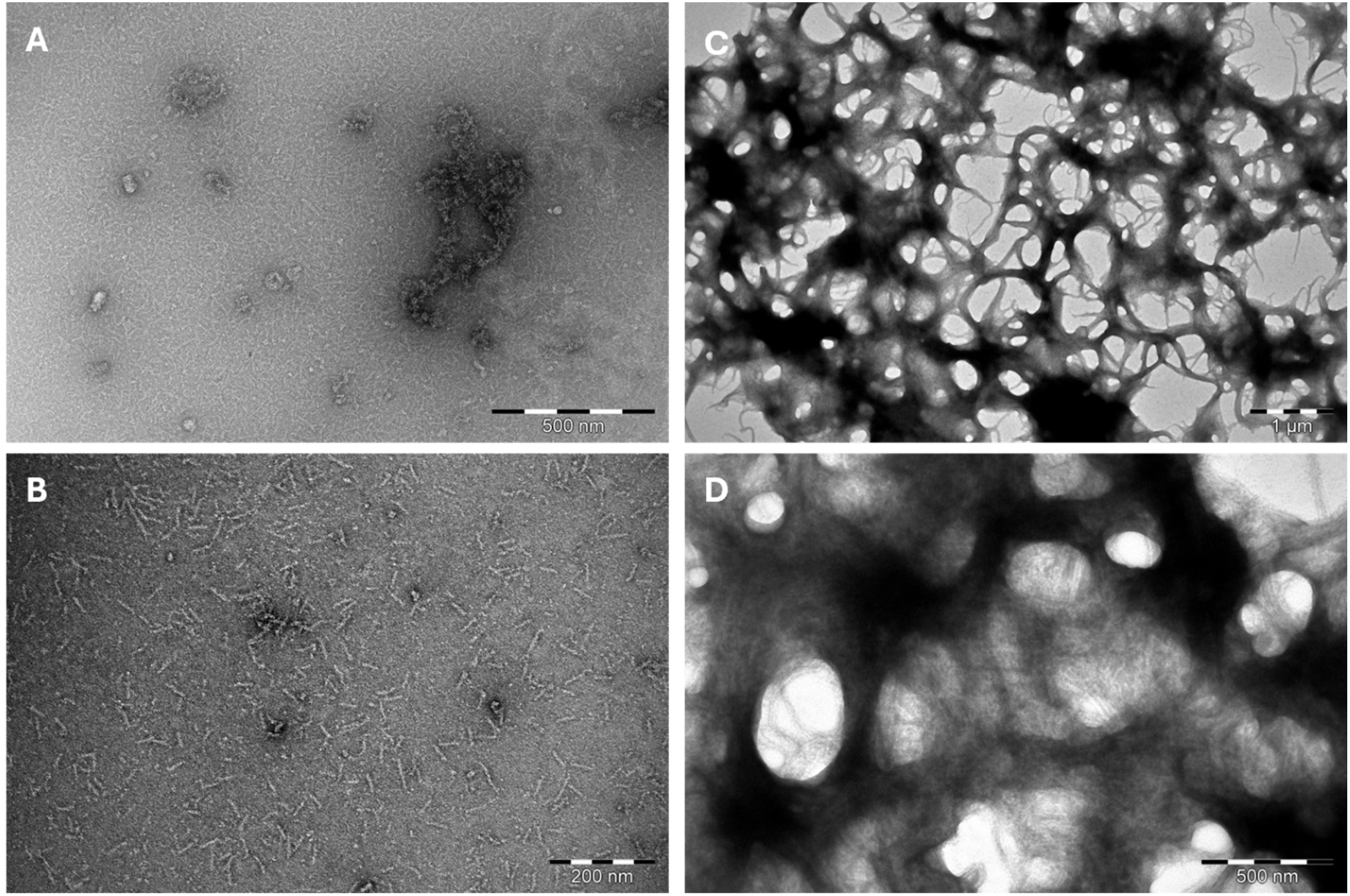
Aggregation of human septins upon the addition of UR214-9. Septin at 0.05 mg.mL^-1^ were incubated with (C, D) or without (A, B) UR214-9 10 µM in 50 mM Tris-HCl pH8, KCL 50 mM and images using negative stain electron microscopy. scale bars. A and D: 500 nM, B: 1 µM, C: 200 nM.

### UR214-9 treatment forms straight bundles, rings and nets in septin-2 overexpressing cells

We next examined how UR214-9 (3µM) impacts septin-2 in SKOV-3 cells transiently overexpressing septin-2 using confocal microscopy. Treatment with non-toxic concentration (3µM) of UR214-9 for 48 hours after transiently overexpressing septin-2 using cDNA for 48 hours, resulted in formation of noodles, rings around nucleus and net-like structures with highly thickened septin-2 (Figure-5).

**Figure 5:**
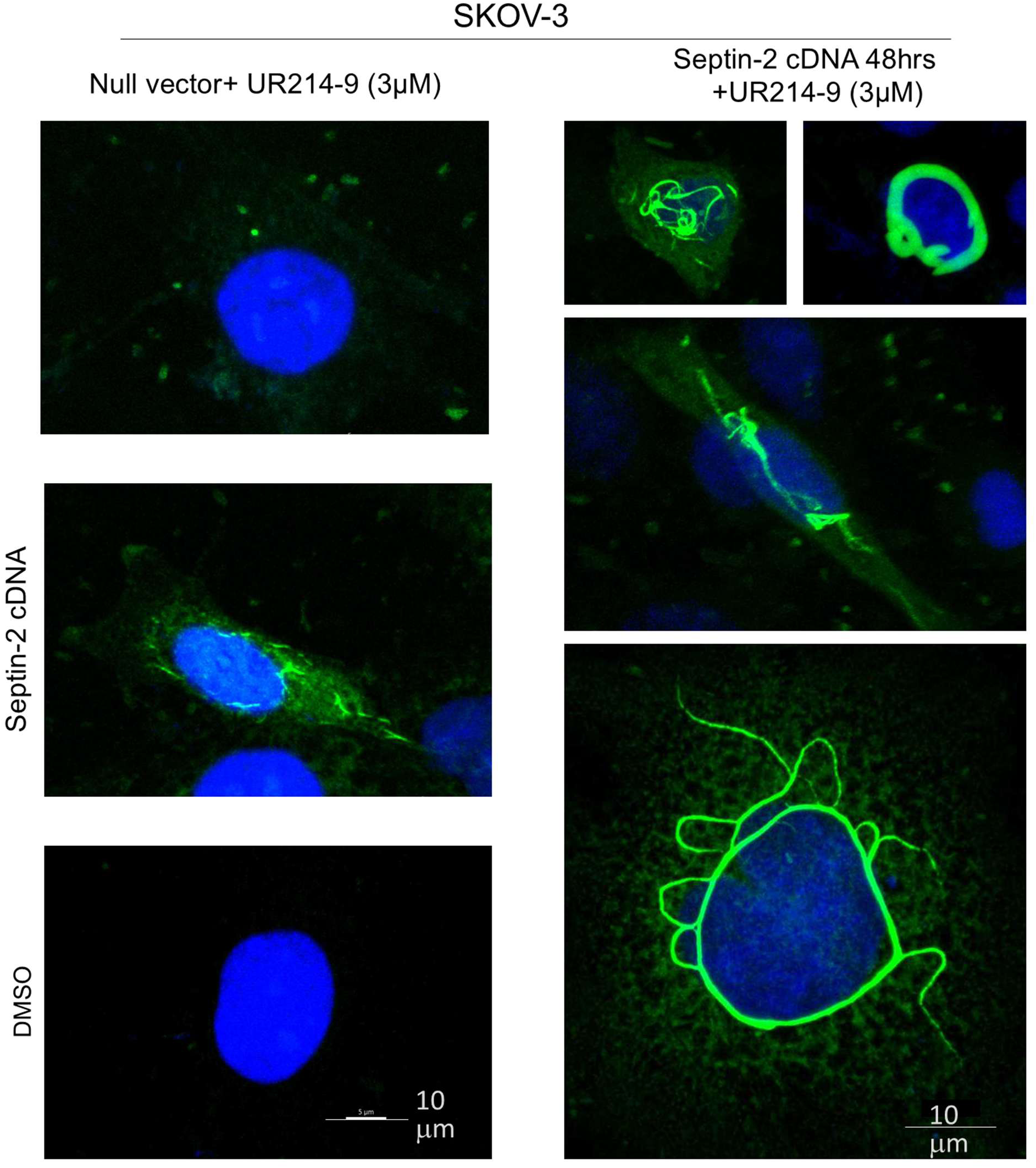
UR214-9 treatment alters septin spatial localization. Treatment with UR214-9 (3µM/48 hours) of SKOV-3 cells transiently overexpressing septin-2 showed bundles of septin-2 filaments which appear in various noodles, rings around the nucleus, and web-like structures. Smaller bar: 5µm. larger Bar=10µm.

### UR214-9 blocks septin-2 localization at cleavage furrow in normal rat kidney fibroblast suppresses cytokinesis in ovarian cancer cells

Septins participate in early cytokinesis events and prime the abscission machinery during cytokinesis^43^. To investigate how UR214-9 impact cytokinesis, we employed a homozygous genome-edited Sept2-EGFP cell line NRK49F-EGFP ^44^, which was devoid of differences between the wild-type and genome edited version during localization of septins over the cell-cycle and normal distribution of septin-2 due to the insertion of EGFP in the start codon in both alleles. The live-cell imaging showed that untreated cells exhibited septin-2-EGFP accumulation at the cytokinesis core during the progression from mitotic pro/metaphase (0 time-point) to anaphase and telophase (50^th^ minute time point) (Figure-6A). Compared to the 0 min time-point when the cells were in metaphase, the cell’s entry in anaphase by the 10^th^ minute timepoint clearly showed septin-2-EGFP localization. During the next 30-40-minute time points, septin-2 rings were seen concentrated at the cleavage furrow. During the next 10 minutes, the septin-2 ring was reduced in size as the cell-division completed. On the other hand, UR214-9(10µM) treated cells showed delayed septin-2-EGP enrichment and such accumulation of septin-2 was witnessed only at the 240^th^ minutes time-point (Figure-6B).

**Figure 6:**
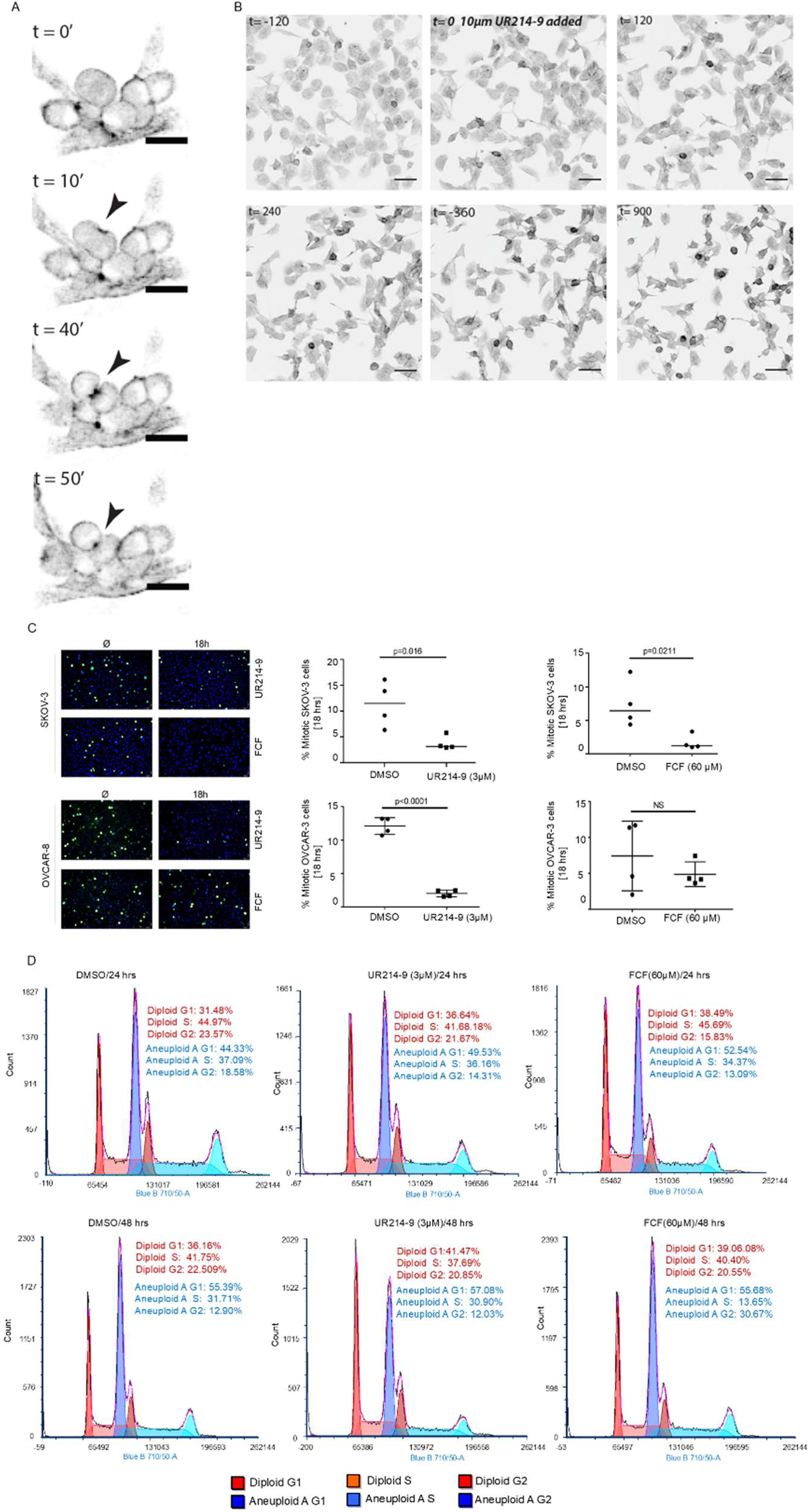
**A):** Live cell imaging of homozygous septin-2 expressing NRK-49F-eGFP cells showed emergence of septin-2 at metaphase (t=0 min). At 10^th^ minute septin-2 is expressed in interphase. At 40^th^ minute timepoint, septin-2 ring is seen concentrated at the cleavage furrow. By the 50^th^ minute time point cytokinesis is nearly completed, and septin-2 levels have reduced. (**B**): On the other hand, UR214-9 (10µM) treatment led to reduced expression of septin-2; cells entering metaphase/interphase and cytokinesis were fewer. (**C**): UR214-9 interferes with mitosis in EOC cell-lines pH3 staining of the vehicle, UR214-9 (3µM) or FCF (60µM) showed that UR214-9 suppresses mitosis in SKOV-3 and OVCAR-8 cells during 18 hours of treatment. FCF also suppresses mitotic cell (pH3) cell population in SKOV-3 cells during 18 hours of treatment. Cells seeded in 8-well glass chamber slides were treated with vehicle, UR214-9 or FCF, fixed, permeabilized and stained with pH3 primary antibody followed by source matched FITC-linked secondary antibody. Four random fields were scoped under a microscope. pH3 positive cells were manually counted and analyzed using GraphPrism. (**D**): SKOV-3 cells (400,000 cells/dish) were seeded, allowed to adhere overnight and then treated for 24 hrs. and 48 hrs. with vehicle or UR214-9 (3µM) in DMSO. Cells were trypsinized, spun down and fixed/permeabilized with cold 70% EtOH. Cells were spun down again, washed with PBST (1 mL) and spun down again. Pellets were suspended in propidium Iodide/RNase solution (Cell Signaling Technology, cat#4087) (300uL), filtered and analyzed using a flow cytometer. Cells were analyzed at each time point separately and ‘forced’ 2 cycle fitting.

**Movie-1:** UR214-9 interferes with cells entering cytokinesis. SKOV-3 cells treated with vehicle or UR214-9 (3µM) and stained with cellMask (FarRed) and SirTubulin (Blue) were monitored using live cell imaging microscope. (**Control**): SKOV-3 cells entering metaphase and progressing to interphase and eventually forming cleavage furrow to complete cytokinesis is shown. (**Treatment**): UR214-9 (3µM) treartment: at 200^th^ minute time-point showed a cell exhibiting cleavage furrow which however showed misalligned daughter chromosomes by 300^th^ minute time point indicating the cytokinesis defects induced by UR214-9.

Similarly, UR214-9 3µM and FCF 60µM suppressed mitotic cell population in SKOV-3 and OVCAR-8 cells significantly (Figure-6C). Phosphohistone-H3 (pH3) antibody detects the condensed chromatin just prior to chromosomal segregation and is considered as a more specific biomarker that stains cells in late-G2 and mitosis, hence stains only mitotically active cells ^45^. PH3 staining led to analysis of cell cycle changes in SKOV-3 cells treated with UR214-9. Both UR214-9 and FCF caused minor G1 phase arrest. FCF treatment at 60µM doses resulted in greater G2 population when counting the diploid population. Among aneuploidic population, similar G1 phase arrest was also seen (Figure-6D) upon UR214-9 or FCF treatment. Extending the treatment for 48 hours retained similar modest G1 phase arrest trend (Figure-6D, lower).

Next, we examined the effects of UR214-9 on cell division in SKOV-3 cells stained with CMG SirTub using live-cell imaging microscopy. SiR-tubulin is a fluorogenic, cell permeable live cell dye that specifically stains microtubules. UR214-9 treatment misaligned tubulins at kinetochore attached spindled microtubules (Movie-1).

### UR214-9 blocks migration, adhesion, SKOV-3 cells and ceramide trafficking without affecting ER & Golgi morphology

Next, we examined the migration of ceramide in SKOV-3 cells by observing the Golgi bodies, as a measure of arising from UR214-9. Reasons for testing Golgi is that septin proteins are required for proper Golgi assembly and function ^46–48^. Secondly, dysregulation of Golgi function is linked to Parkinson’s disease, Alzheimer’s disease and cancer^49^. To determine if UR214-9 treatment disrupted Golgi bodies, live cell staining of SKOV-3 cells was carried out using BODIPY FL C5-Ceramide complexed to BSA (Figure-7A). While compact puncta of labeled Golgi bodies were observed in both control and drug treated cells, additional diffuse ceramide was observed in both UR214-9 and FCF treated cells. Live cells labeled with ER-tracker to observe the endoplasmic reticulum revealed similar staining patterns in all three conditions (Figure-7B), suggesting a Golgi-specific defect in drug-treated cells. Since the ceramide labeling of live cells could be an indirect reflection of trafficking defects or more direct indication of Golgi morphology defects, a batch of cells was treated and fixed to observe Golgi morphology more directly with anti-GM130 staining (Figure-7C). Based on the results of this staining, UR214-9 treated cells exhibited a comparable cis-Golgi morphology to the control cells (Figure-7Cright). In contrast to this result, the FCF treated cells did exhibit changes in the cellular localization of GM130 protein, suggesting that this FCF (not UR214-9) altered the distribution of the Golgi bodies in these cells. Cytokinesis, cell proliferation and migration, invasion and adhesion are each involved in metastasis and tumorigenesis. Depletion of septin-9 in metastatic cancer cells reversed EMT and reduced cell spreading, migration and invasion. In order to test if UR214-9/FCF could alter the migration rate of SKOV-3 cells, the cells were plated on tissue culture plates and then treated for 24 hours with vehicle or UR214-9 (3µM) or FCF (60µM). After treatment, confluent cell monolayers were scratched and then washed to remove free-floating cells. Media with drugs was added back to the cells and images were recorded both 24 and 48 hours after wounding to observe cell migration and wound healing (Figure 7D). Treatment of the cells with concentration of UR214-9 (3μM) inhibited cell migration to an even greater extent than high doses of its parent compound FCF (60μM). Although, both drugs significantly inhibited cell migration compared to the control. UR214-9 treatment blocked adhesion of SKOV-3 cells to fibronectin both at 1 and 3μM (Figure-7E). Similarly, UR214-9 blocked invasion of SKOV-3 significantly at 3μM (Figure-7F).

**Figure-7:**
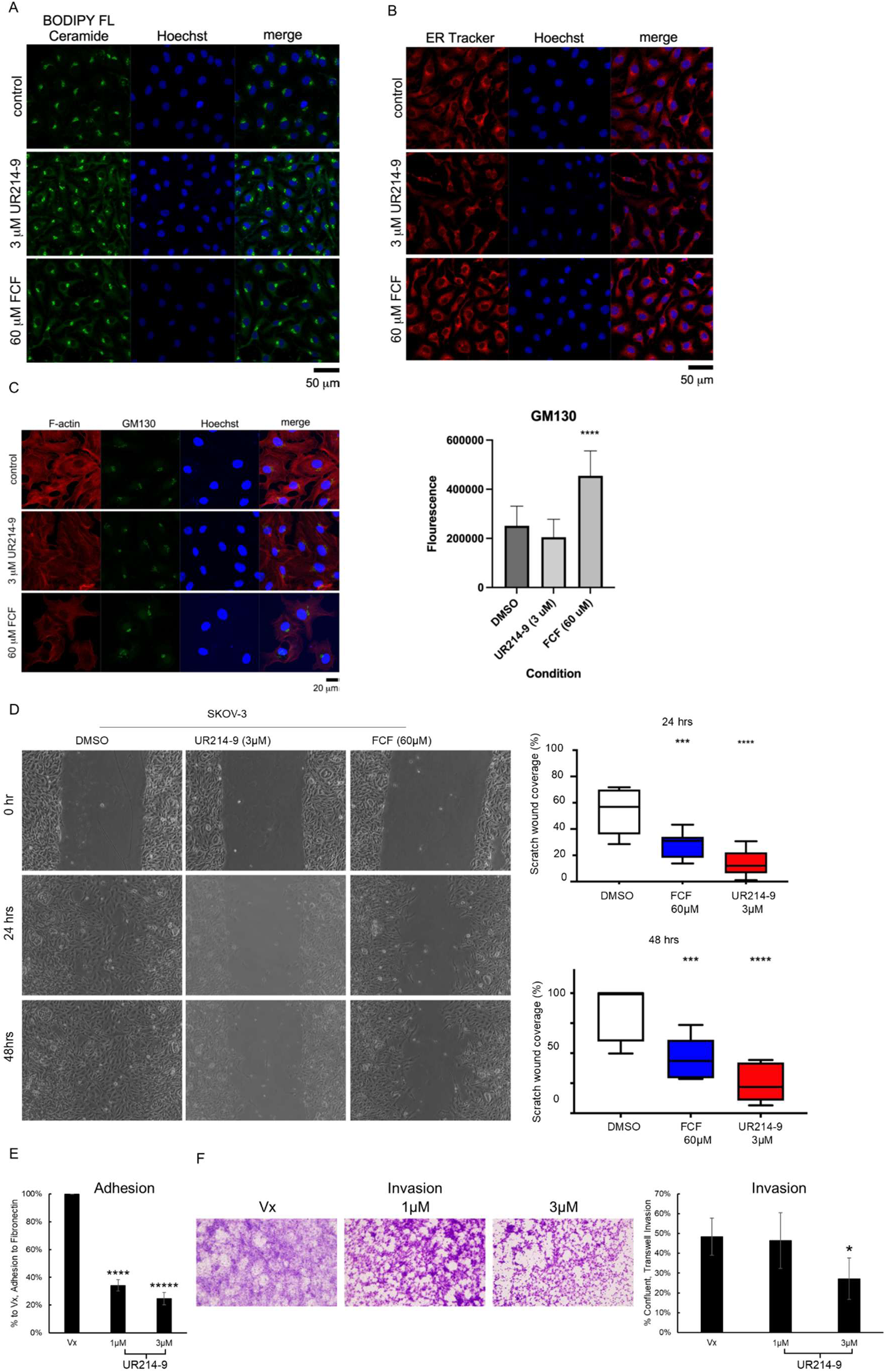
**(A):** UR214-9 and FCF disrupt trafficking of ceramide to Golgi. SKOV-3 cells treated with vehicle, UR214-9 or FCF were imaged after staining with BODIPY-Ceramide. The nucleus was stained with Hoechst 33342 dye. (**B**): UR214-9 does not significantly disrupt ER localization. SKOV-3 cells treated with UR214-9 (3µM) or FCF (60µM) were stained with ER Tracker Red and imaged using a confocal microscope. (**C**-left and right): UR214-9 does not disrupt Golgi morphology (cis-Golgi and localization of GM130) in SKOV-3 cells whereas FCF treatment caused a significant change in localization of GM130 in SKOV-3 cells. (**D**): UR214-9 or FCF treatment blocks cell migration and wound healing in SKOV-3 cells. Adhered cells were pre-treated for 24 hours with vehicle or drug and then scratched. The cells were imaged at 24- and 48-hour post-wounding. Scratched wound coverage area was significantly reduced both in the FCF and UR214-9 treated cells at 24^th^ and 48^th^ hours of observation. (**E**): SKOV-3 cells treated with UR214-9 (1 µM & 3 µM) or DMSO control (Vx) for 24 hours. Both the 1µM (p<0.00005) and 3µM (p<0.00001) had significantly reduced adhesion to fibronectin compared to control. (**F**): SKOV-3 cells treated with UR214-9 (1 µM & 3 µM) or DMSO control (Vx) for 24 hours. The 3µM-treated group had significantly lesser invasion through a Matrigel-coated 5µm-pore Transwell than both the Vx (p=0.0393) and 1µM (p=0.0307) groups

### UR214-9 reduces ovarian cancer cell viability and blocks the growth of ovarian, endometrial and HER2 positive breast cancer xenograft tumors

Treatment with UR214-9 reduced the viability of human ovarian cancer cells (SKOV-3, OVCAR-3, OVCAR-8, and IGROV-1 cells) (Figure-8A). Similarly, UR214-9 inhibited the viability of murine high grade serous ovarian cancer cell-lines which recapitulate fallopian tube origin of ovarian cancer ^50^ (HGS-1 and HGS-3) (Supplementary Figure-6). In addition to the inhibition of growth of ovarian cancer cell-lines *in vitro*, treatment with UR214-9 (25mg/kg, IP, M-F) blocked the growth of SKOV-3 xenograft tumors growing in NSG mice (p=0.03, Figure-8B) without any detrimental effects on the animal weights (Figure-8B-right). Previously we had shown statistically significant expression of septin (2, 6, 7 and -9) in endometrial cancer patients^30^. Therefore, we tested the response of UR214-9(25mg/kg, IP, M-F, once daily) on the growth of AN3CA endometrial cancer cell derived xenografts. UR214-9 treatment under similar doses and treatment methods reduced the growth of AN3CA xenografts in NSG mice than the control on the day-8^th^ and day-17^th^ (p=0.013 and 0.0216 respectively) (Figure-8C). Images of the extracted tumors from the control and treatment groups are shown (Figure-8D). One mouse in the vehicle was found dead during this study and it’s tumor was not extracted. Since, septin-2 was shown to perpetuate HER2 signaling in breast cancer cells^21^, we examined if UR214-9 (25mg/kg, IP, M-F, once daily) treatment can reduce the growth of HER2+ve xenograft. As shown in Figure-8E, UR214-9 treatment resulted in significant reduction in JIMT-1 HER2+ve breast cancer cell-line xenograft. Next, we examined the off-target liabilities stemming from targeting septin/HER2 network in breast cancer cells via examining the global transcriptome changes in JIMT-1 cells, comparing with Afatinib(10nM) versus UR214-9, 1µM, 100X higher concentration of Afatinib) treatment. As shown in Figure-8FG, afatinib, even at low dose of 10nM exhibits massive transcriptomic changes whereas 1µM dose (100x of Afatinib) treatment with UR214-9 showed very limited off-target engagements (volcano plots).

### UR214-9 does not affect vital organs and blood chemistry of healthy C57BL/6 mice

To preliminarily investigate the safety of UR214-9, we treated C57BL/6 mice with vehicle or UR214-9 for 21 days. Peripheral blood collected via mandibular vein puncture when analyzed showed no changes in the complete blood count except monocytes, which portends right for the potentials of UR214-9 tumor treatment because monocytes turn immunosuppressive in malignant settings. Collected organs showed no discoloration or gross damage to the organs (Figure-8H).

## Materials and Methods

### Cell lines, cell culture and reagents

SKOV-3 (ATCC, HTB77), OVCAR-3 (ATCC, HTB-161), IGROV-1 (Sigma, SCC203) and OVCAR-8 (inherited from Laurent Brad’s laboratory) cancer cells were grown in DMEM media (Gibco, 11965) supplemented with 10% fetal calf serum and penicillin (100 units/mL), and streptomycin (100 µg/mL). 2008 (kindly provided by Dr. François X. Claret, University of Texas M.D. Anderson Cancer Center) ovarian cancer cells were grown in RPMI medium (Gibco, 22400) supplemented with 10% fetal calf serum penicillin (100 units/mL), and streptomycin (100 µg/mL). ES2 (ATCC, CRL-1978) was grown in McCoy’s 5A medium (ATCC, 30-2007) supplemented with 10% fetal calf serum penicillin (100 units/mL), and streptomycin (100 µg/mL). Septin-2 (catalog#: HPA018481), septin-7 (catalog#: HPA029524), septin-9 (catalog#: HPA029524) antibodies were purchased from Sigma Aldrich Inc. DyLight 488 (catalog#: DI-1488, rabbit; Dylight (catalog#:DI-2594, mouse) were purchased from Vector Laboratories Inc. Phalloidin-TRITC was purchased from ECM Biosciences (catalog#: PF7551). (−)Epothilone-B (Sigma, catlog#E2556-1mg). Cytochalasin-D (Tocris, catalog#1233). Paclitaxel (Tocris, catalog#1097) were also used in this study.

### shRNA transfection and Western blot analysis of lysates

shRNA plasmid for human septin-2 (Santa Cruz Biotechnology, sc-514206) was transfected to SKOV-3 cells using Lipofectamin 2000 reagent (Thermo Fisher, #116680) following the manufacture’s instruction. Stably transfected cells were selected under the pressure of puromycin (5µg/mL). Individual single cells were cloned by limiting dilution method. Phenotypes of the cloned cells were evaluated by western blotting using anti-Septin-2 antibody (Santa Cruz Biotechnology, sc-514206). For Western blot analysis, septin-2 shRNA-knockdown SKOV-3 cells (clone#9 and -11) along with wild-type and null vector cells were seeded (1-1.5 million) in 100mm^3^ dishes for overnight in complete DMEM media (Gibco, cat#11965-092, supplemented with 10% FBS (Avantar Seradigm, product#89510-186) and 1% Pen-strep (Gibco, cat#15140-122). Media was removed, cells were washed twice with PBS (without Mg++ and Ca++, BioWhittaker, cat#17-512F or Gibco, cat#14190-144). Cells were lysed with lysis buffer (150-200µL, Cell Signaling Tech, cat#9803 supplemented freshly with PMSF). Cells were centrifuged @12000rpm/7 minutes at 4^0^C in cold room, cell debris was removed and clarified supernatant was collected. The protein content was estimated using DC protein assay reagent (A and B) (Bio-Rad, cat#5000113 and 500-0115). 15-20µg of protein lysates were mixed with 12µL/per sample NuPage LDS sample loading buffer mix (75:35 ratio) of (4x, Invitrogen, cat#NP0008, NuPage sample reducing agent, 10x, cat#2270640) and added electrophoresed at 20 volts constant for 90 minutes using NuPage4-12%Bis-Tris gel (Invitrogen, cat#NP0322Box) following manufacturer’s instruments and reagents. Running solution (Invitrogen, NuPage, cat#NP002) was supplemented with NuPage antioxidant solution (Invitrogen, cat#NP0005). After run was complete, proteins were transferred to the PVDF membrane (BioRad, cat#1620177) using filter pads (Bio-Rad, cat#1703966) which were all preincubated in NuPAGE transfer buffer for 10 min before transfer assembly. The transfer was done at room temperature for using semi-dry apparatus using NuPage transfer buffer (Invitrogen, cat#NP0006-1), at 20 V constant voltage for 18 hours. The transfer efficiency was checked with Ponceau staining. Membranes were washed with PBST, blocked with 5% fat free milk for 15-30 minutes and probed with Septin-2 (catalog#: HPA018481), septin-7 (catalog#: HPA029524), septin-9 (catalog#: HPA029524) antibodies (Sigma Aldrich Inc). Secondary HRP-conjugated antibodies were anti-rabbit IgG (1:10,000, Cell Signaling Tech) Chemiluminescent detection was done with a Super Signal West FemtoLuninol/Enhancer (ThermoScientific, cat#1859022 and peroxide cat#1859023) or using Amersham ECL Prime (peroxide solution, cat#RPN2232V2 and enhancer cat#29018903; Cytiva). Next, membranes were stripped using One-minute Western blot stripping buffer (GM Biosciences, cat#GM6001) and reprobed with α-Tubulin antibody and source matched secondary antibody as loading control.

### Synthesis of UR214-9

UR214-9 was synthesized by coupling 3-trifluromethyl,2-fluroaniline with 2,6-dichloro 4-isocyantopyridine in equimolar ration (0.1:0.1) in dry toluene at 85°C overnight. The reaction was monitored using thin-layer chromatography plates with DCM-MeOH or pure ethyl acetate as eluents. Spots were monitored in a UV chamber. The reaction mixture, separated upon completion of the reaction, was filtered under vacuum. The product was washed with hexane, followed by diethyl ether, and was dried under vacuum. A portion of the product was detected in the toluene layer, which was evaporated under reduced pressure and crude residue was purified by thin layer chromatography using ethyl acetate and hexane as eluent. The pure compound band was scrapped off the silica gel and compound were collected by stripping silica gel with Methanol:DCM (9:1). UR214-9 was characterized by X-ray crystallography (Figure-2I). The details of crystallographic data collection, structure solution and refinement and structure description are provided in Supplementary Methods-2.

### Molecular modeling methods

To determine the relative binding of FCF, UR214-9 versus GDP and GTP, we obtained the high-resolution crystal structures of septins from the RCSB ^51^ as targets for molecular docking studies. Specifically, crystal structures of septin isoforms were retrieved, including: septin 2 (PDB ID: 2QNR), septin 6 (PDB ID: 6UPA), septin 7 (PDB ID: 6N0B), and septin 9 (PDB ID: 5CYO). The Swiss-Model^52^ server was employed to model missing atoms and residues in the crystal structures, ensuring structural completeness prior to docking calculations. The modeled and refined structures were subsequently utilized for comprehensive molecular docking investigations. MedusaDock^53–56^, a flexible protein-ligand docking algorithm that accounts for both receptor and ligand flexibility, was implemented for the molecular docking of target compounds, including GDP, GTP, FCF, and UR214-9. The binding pocket for all protein targets was defined based on the co-crystallized ligand positions observed in the respective crystal structures, ensuring biologically relevant binding site selection. To ensure thorough conformational sampling of possible binding modes, each compound was subjected to 1,000 independent docking runs. This extensive sampling protocol allows for comprehensive exploration of the conformational space and identification of energetically favorable binding poses. The final binding poses were evaluated using the Medusa energy function, which incorporates van der Waals interactions, solvation effects, and hydrogen bonding contributions.

### Cell Viability and cell cycle analysis

Cell viability of SKOV-3, OVCAR-3, OVCAR-8, 2008, IGROV-1 and ES2 cells treated with DMSO or varying concentrations of UR214-9 was measured using the Cell Titre96R Aqueous One Solution Cell Proliferation Assay (Promega Corp., catalog#: G3580) following the procedure published earlier^59^.

### Native PAGE and Western blotting of cell lysates

The 100mm3 petri-dishes seeded overnight with 1 million each of the SKOV-3, OVCAR-3 and OVCAR-8 cells,treated with vehicle, UR214-9 (3µM) or FCF(60µM) were placed on ice and the cells were washed twice with PBS (without Ca2+ and Mg2+, before being detached with 200 μl of ice-cold native lysis buffer (80 mM PIPES, pH 6.9, 2 mM MgCl2, 4 mM EGTA, 0.2% saponin, protease+phosphates inhibitor cocktail, Sigma Aldrich). The lysate was collected in 1.5 ml tube and incubated on ice for 10 min. The lysates were then centrifuged at 14,000 g for 10 min at 4°C to remove cell debris. To prevent septin polymerization, clarified lysates were supplemented with NaCl, adding 10 μl of NaCl 5 M for each 100 μl of lysate. After 15 min of incubation on ice, the lysates were clarified in a second centrifugation step of 10 min, 14,000 g at 4°C. Protein quantification was done using the Bradford Protein Assay, and the remaining clarified lysates were kept at −20°C until Native PAGE analysis. The lysates were analyzed by 4–16% Native PAGE using precast Bis-Tris Mini Protein Protein Gels (BN1003BOX; Invitrogen) following the manufacturer’s instructions. The molecular mass marker was NativeMark Unstained Protein Standard (LC0725; Invitrogen). For the Western blot, the gel, the PVDF membrane (BioRad, cat#1620177) and filter pads (Bio-Rad, cat#1703966) were all incubated in NuPAGE transfer buffer for 10 min before assembly. The transfer was done at 4°C for 20 h at 20 V constant voltage. The transfer efficiency was checked by destaining the membrane with an aqueous solution containing MeOH (25%) and acetic acid (10%). The protein marker was identified, and the membranes were completely destained with pure methanol for 3-5 min. The membrane was then blocked with 5% fat free milk and stained with Septin-2 (catalog#: HPA018481), septin-7 (catalog#: HPA029524), septin-9 (catalog#: HPA029524) antibodies (each purchased from Sigma Aldrich Inc). Secondary HRP-conjugated antibodies were anti-rabbit IgG (1:10,000, Cell Signaling Tech) Chemiluminescent detection was done with a Super Signal West FemtoLuninol/Enhancer (ThermoScientific, cat#1859022 and peroxide cat#1859023) or using Amersham ECL Prime (peroxide solution, cat#RPN2232V2 and enhancer cat#29018903; Cytiva).

### Histochemical analysis of tumor tissues cells and co-localization protocol

To determine the impact of UR214-9 treatment on Septin-2 structure in cells, SKOV-3 cells were seeded on glass slides and allowed to adhere overnight. The media was replaced with complete DMEM media supplemented with DMSO or UR214-9 (1µM and 70nM) and cells were incubated for 48 hours. Media was replaced again with new complete medium and fixed with neutral buffered formalin for 15 minutes at 4^0^C. Media was removed, and cells were washed repeatedly with PBST (5x 5mL). The cells were stained with Septin-2 antibody (Sigma Aldrich, catalog #HPA018481) in PSB overnight at 4^0^C. Media was removed again, and cells were washed with 2×5mL PBST. The cells were stained with fluorescence-linked secondary antibody for 1hr. Slides were washed repeatedly in dark for 7×5mL PBST, mounting medium containing DAPI (Vector labs) were applied and covered with glass slide. The slides were stored in dark at 4^0^C till analysis. Confocal images were obtained and processed essentially as published earlier^60^. Confocal images were acquired with Nikon C1si confocal microscope (Nikon Inc. Mellville NY.) using diode lasers 402, 488 and 561. Serial optical sections were obtained with EZ-C1 computer software (Nikon Inc. Mellville, NY). Z series sections were collected at 0.3µm with a 40x PlanApo lens and a scan zoom of 2 or with a 60x Plan Apo objective and a scan zoom of 2, collected every 0.25 µm. Deconvolution measurements were performed with Elements (Nikon Inc. Mellville, NY) computer software. Five cells were outlined and analyzed per field. Actin (stained with phalloidin) Images from fixed cells were acquired on a Zeiss LSM 710 laser scanning confocal microscope with a 63x objective and Zen imaging software suite.

### Ovarian tumor microarray imaging

Twelve bit gray-scale images were acquired with a Nikon E800 microscope (Nikon Inc. Mellville NY) using a 40x Plan Apo objective. A Spot RT3 digital camera (Diagnostic Instruments, Sterling Heights MI) was used to acquire the images. The cameras built-in green filter were used to increase image contrast. Camera settings were based on the brightest slide. All subsequent images were acquired with the same settings.

### Transmission Electron Microscopy

Human octameric septins (Sept2-Sept6-Sept7-Sept9-Sept9-Sept7-Sept6-Sept2) were co-expressed and purified as described previously^61^ Iv et al. (doi: 10.1242/jcs.258484). Septins at 0.05 mg.mL-1 were incubated with UR214-9 at 10 µM for 30 minutes in a 50 mM Tris-HCl, pH 8, KCl 50 mM buffer. 4 µL of the solution was deposited on a glow-discharged carbon-coated electron microscopy grid (CF300-Cu, Electron Microscopy Sciences) and left for incubation for one minute. Grids were stained with a solution of 2% uranyl acetate and air-dried prior to TEM observation. TEM images were acquired using a Tecnai Spirit (Thermofischer, FEI) microscope operated at 80 kV and equipped with a charged couple device camera (Quemesa, Eloise).

### Cell cycle analysis

500,000 cells each of SKOV-3 wt and scrambled shRNA or septin-2 shRNA partial knockdown clones (#9 and 11) were seeded and allowed to adhere in 100mm^3^ petri-dishes for 24 hours. The cells were gently scrapped, centrifuged (1200rpm, 5 minutes) and media was removed. Cells were washed with PBS (1mL), centrifuged again at 1200rpm (3 minutes). Media was removed again, and cells were treated with cold 1mL 70% ethanol for 10 minutes. The cells were centrifuged again at 1200 rpm and ethanol was removed. The cell pellets were transferred to centrifuge tubes and treated with PI/RNase staining solution (300µL, Cell Signaling, cat#4087S) in dark for 30 minutes. The cell suspension was filtered to remove clumps and analyzed by flow cytometry.

### Live cell imaging

SKOV-3 cells (20,000) were seeded in Ibidi (µ-Slides, 4-well, cat#80426) in complete DMEM media and allowed to adhere. Media was replaced with vehicle or UR214-9 (3µM) containing complete DMEM media. Cells were stained with Cell Mask (Thermofisher cat#C37608I) (1:1000 dilution in complete DMEM media), then added 20ul of this to 200ul media in the well. Microscope details and settings include: Leica SP5 inverted point scanning confocal, Lens: 40x 1.25 oil, Resolution:1.3187 pixels per micron, Frame interval: 600 s.

### Wound healing assay

Cells were plated in 6 well tissue culture plates and allowed to adhere overnight before adding drugs. Confluent cells were pretreated for 24 hours with vehicle or drug and then scratched. Three rinses were used to remove loose cells and then media (containing vehicle or drug) was replaced. Cells were imaged 24 and 48 hours after wounding with a 10x objective on a Zeiss Axio Vert A1 inverted microscope and Axiocam 105 color camera. Wound areas were calculated using ImageJ software.

### Cell Adhesion assay

SKOV-3 cells were plated in 6-well tissue culture plates and allowed to adhere overnight before treatment with drugs. Cells at ∼80% cell density were treated pretreated for 24 hours with vehicle or drug and then trypsinized, washed twice in serum-free media, and then seeded at 2,000 cells/well on 96-well plates coated with 50μg/mL fibronectin (Santa Cruz, Cat# sc-29011). Cells were allowed to adhere for 30 minutes, then gently washed to remove nonadherent cells. Adhered cells were fixed with 10% TCA and analyzed using SRB to measure cell density.

### Transwell Invasion assay

SKOV-3 cells were plated in 6-well tissue culture plates and allowed to adhere overnight before treatment with drugs. Cells at ∼80% cell density were treated pretreated for 24 hours in serum-free media [migration buffer] with vehicle or drug, trypsinized, washed twice in serum-free media, and then seeded at 5,000 cells/well in 5μm pore-sized transwells (Corning, Cat#: 3421) coated in 75μL of 1:1 diluted Matrigel (Corning, Cat#: 356255) in sterile water. Migration buffer with 20% FBS added to bottom of well as chemoattractant and incubated for 24 hours at 37°C, 5% CO2. Matrigel and unmigrated cells removed the next day with sterile cotton applicator and washed in sterile PBS. Bottom of transwell with adherent migrated cells fixed in 70% EtOH for 15 minutes, then washed again in PBS. Cells were stained in 0.2% (w/v) crystal violet for 5 minutes, then washed in PBS to remove extra dye. Cells were imaged on an Olympus BX41 using CellSens software, and images were analyzed using ImageJ to measure the density of migrated cells.

### Live Golgi and ER Stains

For live cell staining, cells were stained with BODIPY FL C5-Ceramide complexed to BSA (Invitrogen Molecular Probes catalog# B22650) or ER-Tracker™ Red (BODIPY™ TR Glibenclamide) (Invitrogen Molecular Probes catalog# E34250) and Hoechst 33342 (Invitrogen Molecular Probes) dye according to the manufacturer’s procedure. For Golgi staining, cells were incubated with 5 µM ceramide in HBSS buffer for 30min at 4°C, then rinsed and incubated with fresh buffer containing Hoechst stain for at 37°C for 30 min before being rinsed again and imaged live. For ER staining, cells were incubated with ER-Tracker and Hoechst stains in HBSS buffer for 30 min at 37°C, then rinsed and imaged live. Live cell images were acquired with Zeiss LSM 710 laser scanning confocal microscope with a 20x objective and Zen imaging software.

### Golgi morphology

For viewing Golgi morphology in fixed cells, immunofluorescent staining was carried out using mouse anti-GM130 antibody (BD Transduction Laboratories cat# 610823). Images from fixed cells were acquired on a Zeiss LSM 710 laser scanning confocal microscope with a 63x objective and Zen imaging software suite. Integrated intensity for each cell of fluorescent stain was quantified using ImageJ software and graphed in Prism8.

### Xenograft studies to evaluate antitumor response of UR214-9

All animal experiments were performed only after IACUC approval (2016-011E) from University of Rochester Committee on Animal Resources (UCAR) was obtained. NSG mice were implanted in their left flank with 1 million SKOV-3 cells (volume injected: 100µL; number of animals=10) cells each in matrigel:DMEM (serum free) media (1:1). After 7 days mice were tagged and subdivided into vehicle and treatment groups. SKOV-3 formed palpable tumors within a week and were treated with vehicle or UR214-9 (25mg/kg, IP, M-F). The vehicle formulation was: 40% Hydroxypropyl-beta-cyclodextrin [Acros Organics] & solutol HS15 (Sigma] in sterile water). Similarly, AN3CA cells (100,000 cells/animals were implanted subcutaneously in NSG mice. Nine days post implantation, 25mg/kg equivalent of UR214-9 (1µL=200µg in DMSO) was dissolved in 600uL PBS+400uL of the vehicle and vortexed to obtain a clear suspension. A clear solution was not obtained, and drug was injected with IP as suspension. In addition, JIMT1 and PANC-1 cancer cells were injected into NSG mice which were treated with UR214-9 at a similar dose/route of administration. Tumor burden and animal weights were measured manually by digital calipers on weekly routine. Tumor volume was calculated using the formula ½(L x W^2) where L is a longest diameter and W represents widest width. Statistical difference between the vehicle and treatment groups was analyzed by GraphPrism-8 software using one-way Anova analysis. p<0.05 was considered significant. Mice after the treatment period were euthanized and tumors were resected and snap frozen in liquid nitrogen.

### Preliminary toxicity study in non-tumored rodent model

Healthy female C57BL/6 mice were divided in vehicle(n=5) and UR214-9(n=5) (25mg/kg) group and treated Monday-Friday IP/once-daily. The animal weights were recorded periodically and overall health, fur, socialization behaviors were monitored daily. On the 21^st^ day, 25-40µL blood was collected via mandibular vein puncture from each mouse and stored in EDTA coated blood tubes. Mice were euthanized under CO2 and vital organs were collected, weighed and imaged using iPhone-15. The complete blood count (CBC) was analyzed using the HESKA Element HT5+ instrument. The difference in hemoglobin HB and cell-type population was analyzed using GraphPrism-8 software. One-way Anova analysis was performed. p<0.05 was considered significant.

### Data acquisition and statistical analysis

The prognostic assessment of septin-2/7/9 in the panel of different cancers was conducted using the data available at Human Protein Atlas tools and https://r2.amc.nl/. Microarray databases were analyzed by their tools to determine the impact of septins enrichment on the survival prospects of patients diagnosed with ovarian, liver, pancreatic, lung and breast malignancies. p<0.05 were considered significant. In the cell viabilities one tailed pair distribution T-test was performed. For xenograft studies, a repeated measures analysis of variance was performed using maximum likelihood estimation with group, day, and the interaction between group and day as fixed effects. The correlation of repeated measures on the same subject over time was handled using an unstructured covariance which was allowed to vary by treatment condition. Model assumptions were verified graphically. Analysis was conducted using SAS v9.4 Proc Mixed (Cary, NC). As a sensitivity analysis, the AUC was used to summarize each tumor’s volume over time. The treatment groups were compared with respect to tumor volume AUC using a nonparametric Wilcoxon Rank Sum Test.

## Discussion

Septins are GTP-binding proteins that assemble into intriguing higher-order oligomeric structures and shapes. Such complex septin architectures are typically absent in normal cells, and the mechanisms underlying their emergence and functional tolerance in resource-limited cancer cells remain unclear. Here, we establish a hierarchically dominant role for septin-2 over septin-7 and septin-9. shRNA-mediated depletion of septin-2 reduces septin-7/9 levels, decreases the DNA doublet population, suppresses tumor growth, and enhances sensitivity to microtubule-targeting agents such as paclitaxel and epothilones.

Currently, only three septin-targeting agents:FCF, REM127, and UR214-9, have been reported. In contrast to forchlorfenuron (FCF) and REM127, which promote septin polymerization, UR214-9 disrupts incorporation of septin-2/7/9 subunits into canonical hetero-octamers. This leads to relocalization of septin-2 and septin-9 to the plasma membrane and failure to concentrate at the cleavage furrow, a process essential for cytokinesis.

We further demonstrate that UR214-9 drives the formation of large septin aggregates and higher-order structures. It converts co-expressed human septin octamers (Sept2–Sept6–Sept7–Sept9–Sept9–Sept7–Sept6–Sept2) into large aggregates and induces septin-2-rich filaments, rings, and network-like structures encircling the nucleus upon transient overexpression. This approach enables experimental reconstruction of previously enigmatic septin architecture observed in cancer cells in-vitro.

Functionally, these aggregates likely impair septin-2 redistribution during the interphase-to-cleavage furrow transition, as observed in NRK-49F-SEPT2-EGFP cells, and disrupt cytokinesis in SKOV-3 cells. Consequently, cell proliferation, adhesion, invasion, and migration are inhibited, while essential processes such as ceramide transport to the Golgi and ER–cis-Golgi integrity remain preserved. These effects translate into reduced tumor growth in ovarian, endometrial, and breast cancer xenografts, with minimal evidence of off-target engagement based on global transcriptomic analyses of JIMT1 and PANC-1 cells.

Septins are largely understudied cytoskeletal elements, and therapeutic development has remained extremely limited, with the notable exception of the clinically tested agent REM127, due to several unresolved questions. It is unclear which septin subtype(s) should be prioritized for therapeutic targeting and whether effective strategies will require subtype-selective or pan-septin agents. The septin field also lacks robust, scalable high-throughput reporter assays capable of screening large and chemically diverse libraries with subtype specificity. In addition, the optimal mode of pharmacologic intervention remains uncertain, including whether agents should inhibit septin GTPase activity or disrupt higher-order oligomerization and filament assembly. This uncertainty complicates interpretation of pharmacologic outcomes in the context of septin structural organization (rings and cages), spatial assembly (cortical or submembrane localization), hexa–octameric polymerization, and downstream signaling outputs, including MAPK activation, growth factor signaling, kinase cascades, and secretory regulation.

In this study, we identify septin-2 as a dominant therapeutic target in EOC. Septin-2 is overexpressed in both tumor and stromal compartments and is associated with poor prognoses (Figure-1A; Supplementary Figure-1B). Septin-2-knockdown suppresses tumor growth, reduces septin-7 and septin-9 levels (Figure-1D), both of which independently correlate with poor prognosis in EOC (Figure-1A) and affects cytokinesis and enhances responses to paclitaxel and epothilone-B (Figure-1E, 1G; Supplementary Figure-3).

Next, we investigate whether targeting GTPase activity or targeting oligomerization is the optimal pharmacologic strategy to block septin-driven oncogenesis. First, GTPase activity is not essential for septin filament formation, which regulates cell shape, polarity, cytokinesis, migration, vesicle trafficking, and receptor signaling^62^. Further, since septin-6 lacks GTPase activity and REM127 exerts its anti-septin activity via acting on septin-6, and because septins hydrolyze GTP very slowly, targeting polymerization than GTPase function can provide an effective therapeutic approach in malignancies^63^. This is supported by clinical success of paclitaxel and epothilone-B. Tubulins are functional GTPase like septins. While therapeutic strategies in septins and tubulins can benefit by focusing on septin polymerization, GTPase inhibition has also emerged as a successful clinical strategy for cancer treatment recently, focusing on inhibiting small GTPases (Ras, Rho, Rab) which were often considered “undruggable”^64^.

Next barrier in targeting septins is the lack of structurally diverse chemotypes that interact with septins. To date, other than UR214-9, FCF (Figure-1H) and REM127/REM392^29^ remain the only chemical ‘hits’ and ‘clinical leads’ that interfere with septin. REM127 was also evaluated in phase-ii clinical trials for treatment of Alzheimer’s disease. Functionally, FCF, REM127 and UR214-9 are different from each other, even though UR214-9 is an analog of FCF. While FCF stabilizes the septin-9 octameric oligomers (Figure-3A, right) and REM127 promotes septin polymerization, UR214-9 uniquely prevents incorporation of septin-2/7/9 in the octamer units (Figure-3A-middle). Similarly, while REM127 binds to septin-6 and FCF shows greatest binding to septin-2, UR214-9 shows greatest binding to septin-9 followed by septin-2 (Figure-2B). Further, while REM127 activates calcium signaling, FCF and UR214-9 cast no effects on calcium signaling per se, rather a pretreatment with FCF was found to inhibit Thapsigargin induced Ca^++^ increase (unpublished data). REM127 characterized as septin6/7 molecular glue, carries the problematic accumulation in hepatocytes^65^, and off-target effects and high-doses required for the biological effects, UR214-9 shows limited off-target transcriptomic engagements (Figure-8H). These findings demonstrate that combined in-silico docking, ΔG (kcal/mol), and IDF scoring provide a semi-high-throughput strategy for discovering next-generation UR214-9 analogs or entirely new septin-interacting chemotypes. Following the in-silico screening and hits identification, the activity of the ‘hits’ can be evaluated by the native gel-based immunoblotting assay (Figure-3A) to provide a quantitative and confirmatory estimate of the activity of compounds on the septin polymerization. Next, we address how to read the pharmacologic effects of a septin targeting of FCF/UR214-9 class. The inhibition of septin-2/7/9 incorporation into octamers by UR214-9 casts multi-dimensional impacts on state of septin polymers. Within the cells, septins-2/9 translocate from the cortex to cells periphery as bundles (Figure-3B) and blocks septin-actin co-localization in multiple EOC cells including SKOV-3 (Figure-3C and 3D). The extensive septin polymer-bundling upon treatment of purified septin-2/6/7/8 octamers with UR214-9 seen in cells was also confirmed by electron microscopy (Figure-4). That septins bundle and acquire noodles and random structures wrapped around nucleus when treated with UR241-2 was abundantly shown when we transiently overexpressed Septin-2 in the SKOV-3 cells treated with UR214-9 (Figure-5). Bundled septin-oligomers drastically affect septin’s role in cytokinesis^64^, particularly during the abscission events. Septins are required to complete the abscission but bundled septins fail to be available to complete that process and cytokinesis is blocked. UR214-9 treatment prevents septin-2 from accumulating at the cleavage furrow in NRK-44F-Septin-2eGFP cells (Figure-6A/B). Additionally, α-tubulin is disorganized and unable to form the mitotic spindle in SKOV-3 cells (Movie-1), which results in a reduced pH3 staining mitotic population (Figure-6C). This manifests in G1 phase arrest (Figure-6D). Further, cytokinesis requires membrane trafficking ^65^. Membrane trafficking in cells involves movement of signaling molecules from manufacturing locations in the Golgi to designated release points at the inside of the plasma membrane ^66^. We examined how interfering with septin incorporations in the octamers by UR214-9 may affect Golgi and ER, given that septins are critical for Golgi/ER integrity^67^. Recent studies have associated Golgi function alterations with the metastatic progression of cancer cells. Up-regulation of Golgi-associated genes promotes increased ER-Golgi trafficking and altered secretion and receptor cycling which results in increased cell-matrix adhesion, invasion and migration aiding metastatic progression ^69^. Notably, branched actin filaments facilitate ER/Golgi trafficking^70^. Our study shows that UR214-9 treatment disrupts ceramide trafficking to Golgi (Figure-7A). Interestingly, UR214-9 does not disrupt Golgi morphology as measured by cis-Golgi and localization of GM-130 in SKOV-3 cells (Figure-7C) sparing ER (Figure-7B) at the dose tested. Notably, UR214-9 (3µM) treatment did not affect GM130 localization, but FCF (60µM) treatment led to significant localization of GM130 in Golgi apparatus of SKOV-3 cells (Figure-7C).

Cytokinesis, migration, invasion and adhesion are collaborative processes-^71–72^ and together orchestrate metastasis under the aegis of septins. Interfering with cytokinesis directly impacts migration in SKOV-3 cells (Figure-7D), adhesion (Figure-7E) and invasion (Figure-7F), each of which are complementary elements of metastasis. Interference with cytokinesis by UR214-9 reduced EOC cell-lines cell proliferation of representing serous and clear cell phenotypes *in vitro* during a 48 hrs. treatment duration (Figure-8A). In-vivo, UR214-9 treatment controlled the growth of SKOV-3 ovarian (Figure-8BC) and endometrial cancer xenografts (Figure-8C) and HER2+ve breast cancer growth (Figure-8D) in the NSG mice. Similar reduction in the growth of pancreatic cancer xenograft was observed (data not shown).

Next, we address the concerns regarding off-target liabilities of UR214-9, particularly against hematopoietic and bystander cells. First, RNA-seq analysis in pancreatic PANC-1 and JIMT-1 breast cancer cells (Figure-8F/G) show that UR214-9 treatment affects minimal number of genes^73^. Secondly, targeting septins in bystander cells may not be a significant concern, as many of these cells are tumor-associated and actively contribute to tumor growth, metastasis, and chemoresistance ^74^. Regarding hematotoxicity, UR214-9 is predicted to be less hematotoxic than taxanes, epothilones which indiscriminately destruct hematopoietic cells ^75^. This inference is borne partly from the evidence that hematopoietic cells can function part independently of septin-^76–77^. Second, a pilot studies using non-tumors C57/BL-6 mice. Fifteen (15) intraperitoneal treatments of 25 mg/kg over 21 days did not cause any observable detriment to the well-being of animals or alter hematopoietic cell counts (WBCs, platelets, neutrophils) or hemoglobin levels compared to control male mice (Figure-8E). Xenograft and safety studies were limited by the poor aqueous solubility of UR214-9. As a result, the in vivo efficacy of UR214-9 may be underestimated in these experiments (Figure 8), including effects observed in complete blood analyses. In contrast, in-vitro studies were not affected by solubility constraints. Data described here combined with recently published activities of UR214-9 in metastatic models of breast cancer^78^ and glaucoma^79^ highlight the therapeutic benefits of UR214-9 against diverse diseases. The future goals are to develop an improved delivery formulation and a set of analogs that exhibit greater water solubility.

**Figure 8:**
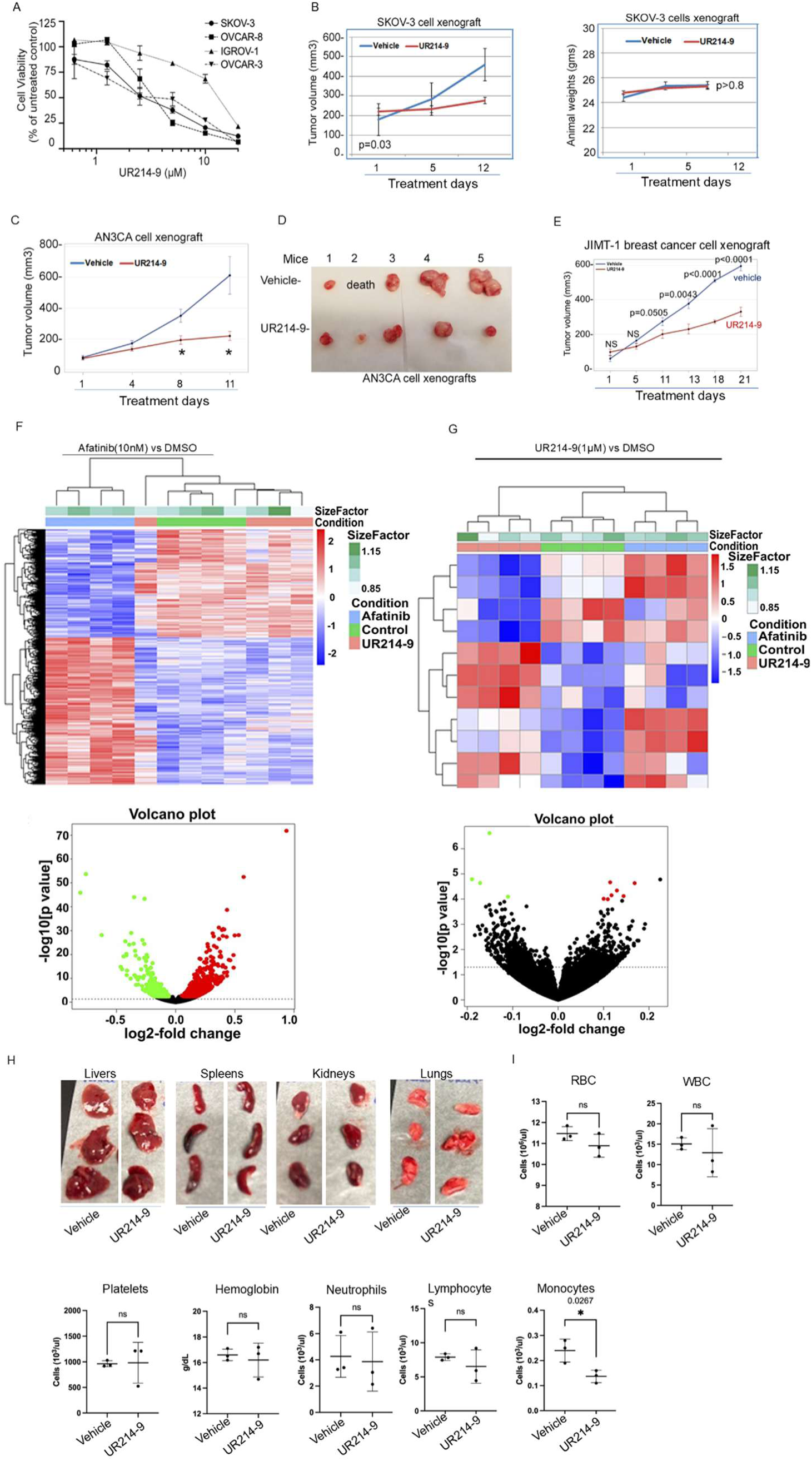
UR214-9 reduces viability of ovarian cancer cell lines and decreases tumor sizes in ovarian, endometrial and HER2+ve breast cancer xenograft experiments. (**A**): UR214-9 treatment suppresses the viability of SKOV-3, OVCAR-3, OVCAR-8, IGROV-1 EOC cells during 48 hours of observation. (**B-**left): UR214-9 (25mg/kg, M-F, IP) treatment blocked the growth of SKOV-3 xenografts in NSG mice during 20 days of observations**. (B-**right): UR214-9 (25mg/kg, M-F, IP) treatment (IP) did not affect animal weights or caused any observable toxicity to animals during the course of treatment. (**C**): UR214-9 (25mg/kg, M-F, IP) treatment blocked the growth of AN3CA endometrial cancer cell derived xenografts in NSG mice during 20 days of observations. Treatment resulted in smaller tumor burden on day-8^th^ and day-17^th^. (**D**): Images of the harvested tumors from Figure-8C mice are shown. (**E**): UR214-9 treatment reduced the growth of JIMT-1 breast cancer xenograft growing in NSG mice. (**F-G**): Hierarchically clustered heat map of mRNA expression for 1234 significantly differentially expressed genes (BH adjusted p-value < 0.05) in the JIMT-1 breast cancer cells treated with afatinib compared to control and associated volcano plot. Hierarchically clustered heat map of mRNA expression for 11 significantly differentially expressed genes (BH adjusted p-value < 0.05) in the JIMT-1 breast cancer cells treated with UR214-9 compared to control and associated volcano plot. Heatmap color key represents row scaling of the rLog transformed expression values. The volcano plots have horizontal lines at p-value 0.05 and individual genes/dots are colored red when the adjusted p-value is >= 0.05 and the fold-change is > 0 and green when the fold-change is < 0. (**H**): To determine, preliminary toxicity of UR214-9 against vital organs and hematopoietic cells, healthy non-tumored C57BL/6 mice were treated with UR241-2 (25mg/kg, IP, once daily) for 21 days. The Complete Blood Count (CBC) of peripheral blood of the C57Bl/6 mice treated with vehicle or UR214-9 collected prior to euthanasia was analyzed by HESKA Element HT5+ instruments. The graphs of the cell-type population in treatment versus vehicle were plotted using GraphPrism V8. one-way ANOVA. p<0.05 was considered significant. Mice were euthanized and organs were collected and imaged.

## Supplementary Figures

**Supplementary Figure-1:** Survival probabilities of liver, renal, pancreatic, lung and melanoma patients exhibiting septin-2, 7 and -9 overexpression.

**Supplementary Figure-2**: original uncut scans of western blot gels of the data shown in Figure-1D and Figure-3A.

**Supplementary Figure-3**: Septin-2kd enhances the responses of the (−) epothilone-B.

**Supplementary Figure-4**: Methods of X-ray crystallographic structural analysis of UR214-9 and images and crystallographic data.

**Supplementary Figure-5:** IFD scores of FCF, UR214-9 and its analogs.

**Supplementary Figure-6:** UR214-9 suppresses the cell viability of high grade serous murine ovarian cancer cell-lines (HGS-1 and -3).

## Author contributions

RKS designed study, conceived and synthesized UR214-9 initially, had it synthesized in contract with Presude Lifesciences Pvt Ltd, and conducted animal experiments with EL, and assembled the manuscript. CWAS designed and performed IHC, recorded and analyzed images, performed adhesion, wound healing, and trans-well assays. MS performed the statistical analysis. NAS performed western blots along with AJ, and NK. NAS also performed the molecular docking initially. CR reexamined survival curves freshly this year using R2-genomics portal. ND performed live cell imaging under supervision of DTB. TK performed the confocal microscopy. KK, AS, SN along with SE and NVD performed different aspects of docking studies. AB contributed the electron microscopy. NY developed knockdown clones. GCE performed migration assay under supervision of JNH. TK and JNH performed different confocal microscopy. JNH also stained the cells for ER and Golgi related studies. TM, NK, JPM, NZ, and KKK conducted western blot, immunoprecipitation, cell viability and other supplementary studies. HE provided NRK-49G-EGFP cells. VH performed confocal microscopic studies. KKK performed the cell viability studies. ET and AB critically examined the manuscript based on their expertise in septins. RGM provided partial resources and edited the manuscript. RBRT provided part resources and edited the manuscript. Every author read the manuscript, participated in editing and approved the final version for submission.

## Ethics statement

Some data used in this study were sourced from publicly accessible databases carrying de-identified patient cases. No patient privacy was violated. Animal experimental study was approved by University Committee on Animal Resources (2016-011E).

## Acknowledgements

RKS acknowledges Mae Goode Foundation award to partially support these studies. RGM provided laboratory and personnel supports. Helge Ewers laboratory provided NRK-49G-EGFP cells. JNH gratefully acknowledges funding support from Colgate University Faculty Research Council. The X-ray instrument at University of Rochester was purchased with funding from NSF MRI program grant CHE-1725028. NVD acknowledges support from the National Institutes of Health (NIH), USA (R35 GM134864) and the National Science Foundation, USA (2040667).

## Material and Data availability

UR214-9 is available on request. A material transfer agreement MTA is required.

